# Genetic risk and inflammatory signaling converge on cell type-specific regulatory programs in type 1 diabetes

**DOI:** 10.64898/2026.09.25.753922

**Authors:** Liu Wang, Zong Wei

## Abstract

Type 1 diabetes (T1D) is a complex autoimmune disease characterized by the destruction of insulin-producing pancreatic β cells. Both genetic susceptibility and epigenetic dysregulation contribute to T1D risk. Recent studies have reported β-cell dysfunction before disease onset, suggesting that β-cell-intrinsic mechanisms contribute to pathogenesis. However, the mechanistic link between disease-associated variants and β-cell dysfunction remains poorly understood. We hypothesized that a subset of T1D-associated variants directly influences cell type-specific candidate cis-regulatory elements (cCREs) by disrupting transcription factor (TF) binding and altering gene expression. To define cell type-specific cCREs in human islets, we integrated publicly available epigenomic datasets, including bulk histone modification profiles and single-cell multiomic data from human pancreatic islets. We combined these data with T1D genome-wide association study (GWAS) variants to identify disease-associated SNPs located within enhancer regions. Using multiomic datasets from PANC-DB, we characterized cell type-specific chromatin accessibility in nondiabetic and T1D islets and identified regulatory regions that may control gene expression in β cells and other islet cell types. We then applied ChromBPNet, a deep learning model that predicts base-pair-resolution chromatin accessibility from scATAC-seq data, to evaluate how specific variants may alter local regulatory activity. In parallel, we used TF footprinting to nominate TFs likely to bind these variant-containing regions. These analyses identified several T1D-associated SNPs predicted to alter chromatin accessibility at candidate TF binding sites, suggesting mechanisms by which noncoding variants may contribute to β-cell dysfunction and T1D susceptibility. Our findings link T1D-associated variants to cell type-specific enhancer activity and regulatory pathways and provide candidates for future mechanistic studies.

## Introduction

Type 1 diabetes (T1D) is an autoimmune disease characterized by immune-mediated destruction of insulin-producing β cells in the pancreatic islets. Genetic susceptibility is a major determinant of T1D risk, and genome-wide association studies (GWAS) have identified numerous loci associated with the disease^1^. A large proportion of T1D-associated variants lie in noncoding regions of the genome, suggesting that altered transcriptional regulation contributes to disease susceptibility^1,2^. However, the functional consequences of many of these variants remain unresolved, particularly in individual endocrine and non-endocrine cell types of the pancreas.

Although immune-mediated β-cell destruction is a prominent feature of T1D, accumulating evidence indicates that β-cell-intrinsic defects also contribute to disease pathogenesis, often at early stages of T1D^3,4^. Changes in β-cell function and identity, cellular stress responses, inflammatory signaling, and interactions with the immune system may occur during disease progression^5,6^. These findings suggest that β cells are not simply passive targets of immune-mediated destruction, and they raise the possibility that genetic variation affecting β-cell regulatory programs contributes to cellular dysfunction and disease susceptibility. Defining how T1D-associated variants influence regulatory mechanisms within β cells and other non-immune cell types is therefore important for understanding the molecular processes connecting genetic risk to disease pathogenesis.

Noncoding genetic variants can influence gene expression by altering the activity of candidate cis-regulatory elements (cCREs), including enhancers and promoters^7,8^. cCRE activity is associated with characteristic epigenomic features^9,10^, including chromatin accessibility and histone modifications such as H3K4me1 and H3K27ac. Integration of disease-associated variants with these regulatory annotations can therefore help identify candidate variants with potential regulatory functions. Pancreatic islets are composed of multiple endocrine and non-endocrine cell populations with distinct transcriptional and epigenetic programs. Regulatory signals measured in bulk islets may therefore obscure effects that occur only within specific cell types.

Single-cell and single-nucleus gene expression profiling can now resolve the regulatory landscapes of individual cell types and states^11,12^. Single-cell assay for transposase-accessible chromatin using sequencing (scATAC-seq) and single-cell multiomic approaches have identified cCREs in individual cell populations. Utilizing these platforms, large-scale human islet consortiums (IIDP, HPAP, and PanKbase) have generated genomic, epigenomic, and transcriptomic datasets from pancreatic tissue of donors with and without diabetes^13–17^. Integrating these datasets with T1D GWAS variants enables identification of disease-associated variants located in cCREs and of the pancreatic cell types in which these regions may function.

However, the presence of a disease-associated variant within a cCRE does not establish that the nucleotide substitution itself alters regulatory activity. Sequence variation within a cCRE may modify transcription factor (TF) recognition motifs, TF occupancy, nucleosome organization, or local chromatin accessibility. Sequence-based deep learning approaches have recently been developed to predict these effects directly from genomic sequence^18–20^. ChromBPNet^19^, for example, models base-pair-resolution chromatin accessibility while accounting for sequence biases in chromatin accessibility assays, enabling prediction of the regulatory consequences of sequence alterations. Complementary approaches for TF footprinting and single-cell regulatory analysis can further identify candidate TFs whose occupancy may be affected at variant-containing regulatory regions. Combining these approaches can move the analysis from annotation of disease-associated variants toward mechanistic predictions of their regulatory effects. Here, we integrated T1D-associated genetic variants with human pancreatic islet epigenomic and single-cell multiomic datasets to characterize candidate regulatory variants across pancreatic cell types. We combined enhancer-associated histone modifications with chromatin accessibility profiles to define regulatory regions and identify T1D-associated variants located within these elements. We used single-cell chromatin accessibility data to determine the cell type specificity of variant-containing regulatory regions and to examine their regulatory landscapes in control and T1D islets. We then applied sequence-based deep learning to evaluate allele-dependent effects of selected variants on local chromatin accessibility and integrated these predictions with TF footprinting to identify candidate TFs whose binding may be altered by T1D-associated sequence variation. Together, these analyses connect T1D GWAS variants with cell type-specific cCRE activity and nominate candidate noncoding variants and regulatory mechanisms.

## Results

### Single-cell multiomic profiling defines the regulatory landscape of human pancreatic islets in T1D

To characterize the cellular and regulatory landscape associated with T1D, we analyzed publicly available single-cell multiomic data from PANC-DB (Human Pancreas Analysis Program [HPAP])^13,21–23^, comprising pancreatic islet samples from 20 donors: 10 non-diabetic (ND), 2 autoantibody-positive (AAb+), and 8 T1D donors (Fig. 1A; Supplementary Table 1). After quality control and integration, we retained 67,734 cells, including 37,627 ND, 7,034 AAb+, and 23,073 T1D cells. UMAP visualization of the scRNA-seq profiles resolved the major endocrine and non-endocrine pancreatic cell populations (Fig. 1B), and the scATAC-seq profiles showed corresponding cell type-specific chromatin accessibility landscapes (Fig. 1C). Joint analysis of gene expression and chromatin accessibility identified nine major pancreatic cell populations across disease groups (Fig. 1D), comprising endocrine α, β, δ, and PP cells and non-endocrine ductal, acinar, stellate, endothelial, and immune populations.

**Figure 1.**
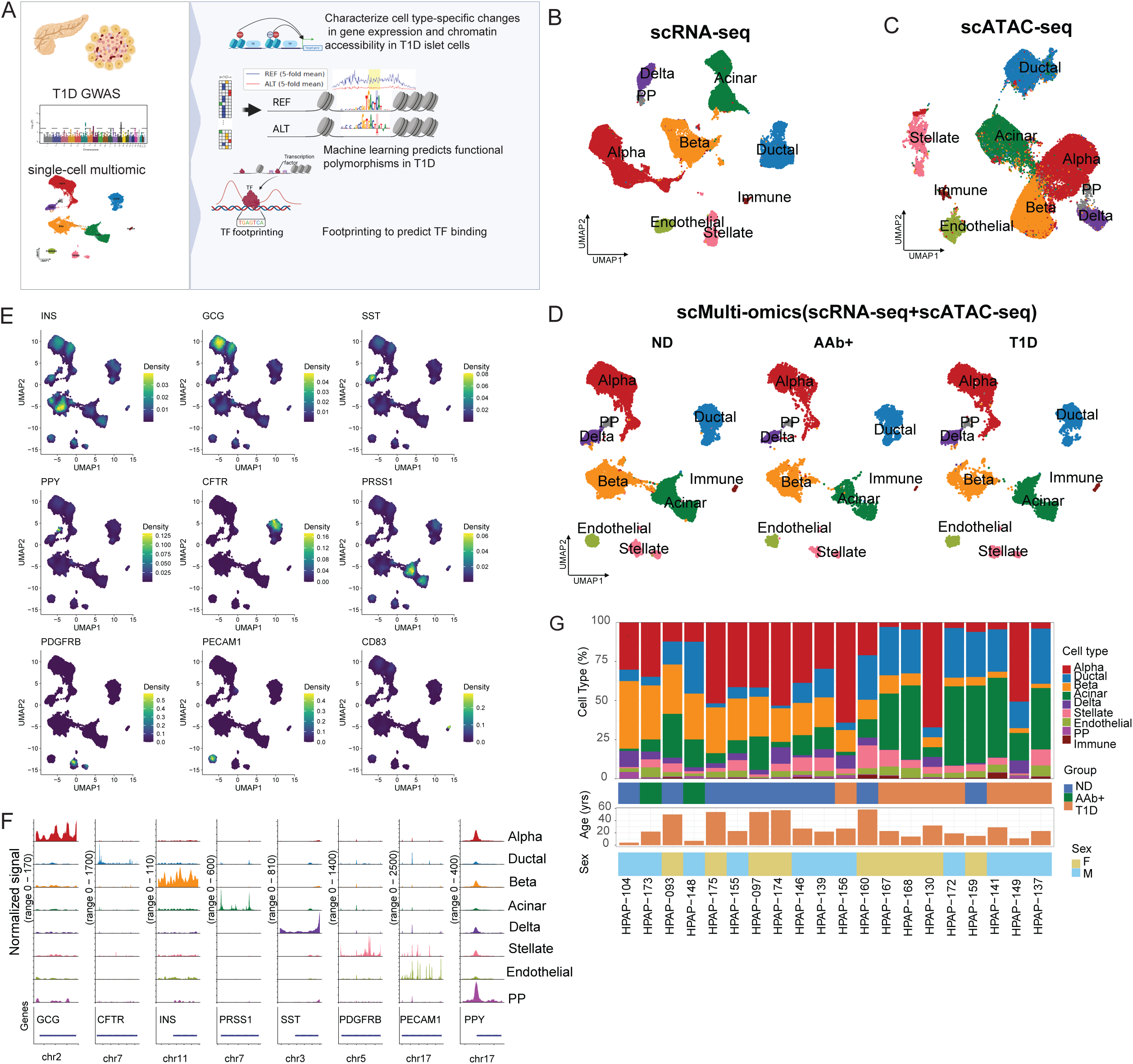
Single-cell multiomic characterization of human pancreatic islets across T1D disease states. (A) Schematic of the study design (created with BioRender.com). Single-cell multiomic data were obtained from PANC-DB for non-diabetic (ND), autoantibody-positive (AAb+), and type 1 diabetes (T1D) donors. (B) UMAP visualization of the scRNA-seq component of the single-cell multiome dataset, colored by annotated cell type, showing the transcriptional organization of major pancreatic endocrine and non-endocrine cell populations. (C) UMAP visualization of the scATAC-seq component of the same single-cell multiome dataset, colored by corresponding cell-type annotations, showing the chromatin-accessibility landscape of major pancreatic cell populations. (D) UMAP showing the clustering of 67,734 single cells by their integrated gene expression and chromatin accessibility profiles. (E) Feature plots showing expression of canonical lineage-marker genes used to support annotation of the major pancreatic cell populations. (F) Normalized aggregate scATAC-seq signal tracks across representative cell-type marker loci, illustrating cell-type-specific chromatin accessibility patterns. (G) Summary of cell-type composition and donor characteristics across samples.

Cell-type annotations were supported by canonical lineage markers, including INS (β), GCG (α), SST (δ), PPY (PP), CFTR (ductal), PRSS1 (acinar), PDGFRB (stellate), PECAM1 (endothelial), and CD83 (immune) (Fig. 1E), together with cell type-specific chromatin accessibility at corresponding lineage-defining loci (Fig. 1F). Of the 67,734 cells, 39,171 (57.8%) were endocrine cells, including 22,107 α (32.6%), 13,391 β (19.8%), 3,009 δ (4.4%), and 664 PP cells (1.0%). The non-endocrine compartment comprised 12,328 acinar (18.2%), 10,217 ductal (15.1%), 3,366 stellate (5.0%), 2,172 endothelial (3.2%), and 480 immune cells (0.7%).

We next examined cell-type composition across individual donors and disease groups (Fig. 1G). The proportion of β cells was markedly lower in T1D islets (7.9% of cells) than in ND islets (24.8% of cells). In AAb+ samples, β cells accounted for 31.5% of cells, although this group comprised only two donors. Overall, endocrine cells represented 65.9% of ND, 58.0% of AAb+, and 44.6% of T1D islet cells, indicating a shift in pancreatic cellular composition in T1D.

### Cell type-specific changes in gene expression and chromatin accessibility across T1D disease states

We first sought to determine which pancreatic cell populations undergo the largest transcriptional and epigenetic changes across T1D disease states. We performed pseudobulk differential gene expression analysis within each annotated cell type and quantified genes increased or decreased in pairwise comparisons of T1D versus ND, T1D versus AAb+, and AAb+ versus ND (Fig. 2A). The extent of transcriptional remodeling varied markedly across cell types and comparisons. Several endocrine and exocrine populations showed large numbers of differentially expressed genes, particularly in comparisons involving T1D, whereas other pancreatic populations showed substantially fewer changes. AAb+ cells were also transcriptionally distinct from ND cells in several populations, suggesting that some molecular changes are detectable before clinical onset of T1D, although the AAb+ group comprised only two donors. Overall, disease-associated transcriptional changes were widespread but highly cell type dependent.

**Figure 2.**
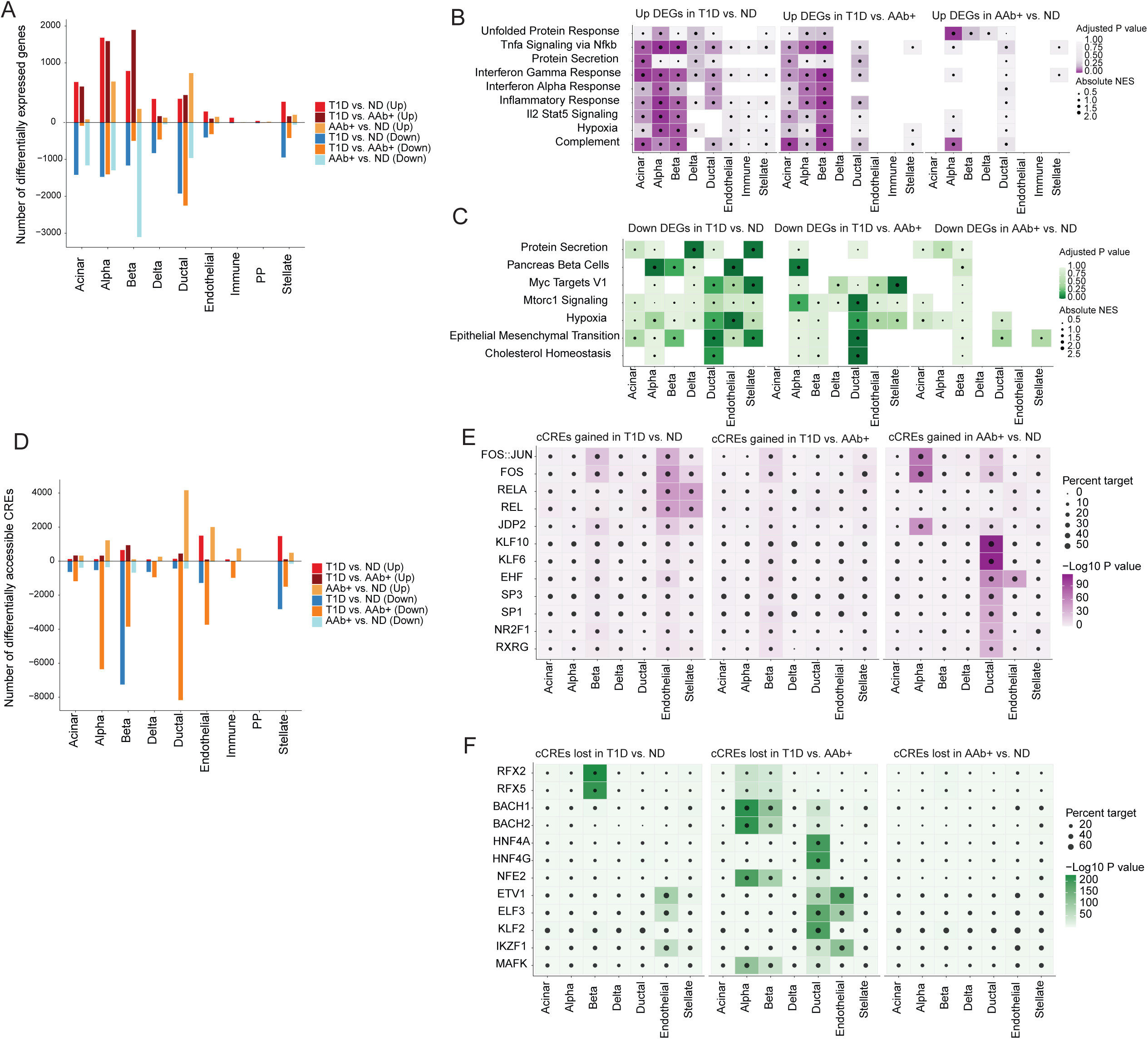
Cell type-specific transcriptional and chromatin regulatory remodeling in human pancreatic islets in T1D. (A) Numbers of significantly upregulated and downregulated differentially expressed genes (DEGs) identified by pseudobulk analysis in each pancreatic islet cell type. (B–C) Preranked gene set enrichment analysis (GSEA) of upregulated (B) and downregulated (C) DEGs across islet cell types, highlighting cell type-specific transcriptional programs associated with T1D. (D) Numbers of differentially accessible candidate cis-regulatory elements (cCREs), including gained and lost cCREs, identified in each islet cell type. (E–F) Transcription factor (TF) motif enrichment analysis of gained (E) and lost (F) cCREs across islet cell types.

We next asked whether these differentially expressed genes converged on common biological pathways. Preranked gene set enrichment analysis^24^ showed significant enrichment of inflammatory pathways among genes increased in T1D (Fig. 2B). Interferon-γ response, interferon-α response, inflammatory response, TNF-α signaling through NF-κB, IL2-STAT5 signaling, hypoxia, and complement were enriched across multiple pancreatic cell populations, with the strength and distribution of enrichment varying among cell types and disease comparisons (Fig. 2B). These inflammatory signatures were most prominent in comparisons involving T1D, consistent with coordinated activation of immune and stress-responsive transcriptional programs in established disease.

In contrast, genes decreased across disease states were enriched for a distinct set of cellular programs (Fig. 2C). These included protein secretion, pancreas β-cell signatures, MYC targets, mTORC1 signaling, hypoxia, epithelial-mesenchymal transition, and cholesterol homeostasis, with substantial variation among individual pancreatic cell populations. The enrichment of inflammatory pathways among upregulated genes, and of secretory, metabolic, and cell identity-associated pathways among downregulated genes, suggests that inflammatory activation in T1D occurs together with disruption of normal pancreatic cellular programs.

Because changes in gene expression can be caused by or correlated with remodeling of the chromatin landscape, we next examined chromatin accessibility across the same disease groups. We performed pseudobulk differential accessibility analysis within each annotated cell type and quantified cCREs whose accessibility was gained or lost in T1D versus ND, T1D versus AAb+, and AAb+ versus ND (Fig. 2D). Extensive chromatin remodeling was detected across pancreatic cell populations, but the number and direction of differentially accessible cCREs differed markedly by cell type and disease comparison. Comparisons involving T1D showed the largest accessibility changes in several populations, with large numbers of both gained and lost cCREs. These findings indicate that the transcriptional remodeling observed across disease states is accompanied by widespread reorganization of the pancreatic cis-regulatory landscape. We next sought to identify TFs associated with these disease-associated accessibility changes. Motif enrichment analysis was performed separately within cCREs that gained or lost accessibility in each cell type and disease comparison (Fig. 2E,F). cCREs gained in disease were enriched for motifs recognized by several inflammatory and signal-responsive TF families, including FOS::JUN, FOS, RELA, REL, JDP2, KLF10, KLF6, EHF, SP3, SPI1, NR2F1, and RXRG (Fig. 2E). AP-1-associated motifs (*FOS::JUN* and *FOS*) and NF-κB-associated motifs (*RELA* and *REL*) were enriched within gained accessible regions in several pancreatic cell populations. Thus, independent analyses of the scRNA-seq and scATAC-seq data converged on inflammatory regulatory programs, supporting coordinated remodeling of transcription and chromatin accessibility in T1D.

In contrast, cCREs that lost accessibility showed a distinct spectrum of motif enrichment, including RFX2, RFX5, BACH1, BACH2, HNF4A, HNF4G, NFE2, ETV1, ELF3, KLF2, IKZF1, and MAFK (Fig. 2F). These motifs showed pronounced cell type-specific patterns across disease comparisons. The enrichment of RFX-family and other lineage-associated TF motifs within lost cCREs suggests that disease-associated chromatin remodeling may involve reduced activity of lineage-defining TFs in endocrine and non-endocrine cells. This interpretation is consistent with the reduction of secretory, metabolic, and cell identity-associated transcriptional programs among downregulated genes (Fig. 2C).

Overall, these analyses reveal extensive and cell type-specific molecular remodeling across T1D disease states. The concordance between inflammatory transcriptional programs and increased accessibility of AP-1- and NF-κB-associated regulatory elements provides complementary evidence that inflammatory signaling is accompanied by remodeling of the pancreatic regulatory landscape. Conversely, loss of accessibility at regulatory elements associated with lineage-associated and other regulatory factors coincided with reduced homeostatic and metabolic transcriptional programs.

### T1D is associated with remodeling of the pancreatic immune compartment and broader inflammatory responses

To define disease-associated changes within the pancreatic immune compartment, we focused on immune cells from ND, AAb+, and T1D donors in the integrated single-cell multiome dataset and asked whether specific inflammatory and lineage-associated programs distinguished T1D from ND immune cells. Dot-plot analysis revealed broad differences across cytokine, myeloid/macrophage, cell-surface, antigen-presentation, and transcriptional-regulatory gene categories (Fig. 3A). Among cytokine and inflammatory-response genes, T1D immune cells showed altered expression of TNF-associated genes, including *TNF* and members of the TNF receptor family, consistent with increased engagement of inflammatory signaling. T1D immune cells also displayed a prominent myeloid/macrophage-associated program, with increased expression and/or a greater fraction of expressing cells for *CD163, C1QA, C1QB, C1QC, SPP1, AIF1, TYROBP,* and *FCER1G*. The coordinated expression of complement-associated macrophage genes together with *SPP1* and *CD163* suggests expansion or activation of a macrophage-like inflammatory state in T1D.

**Figure 3.**
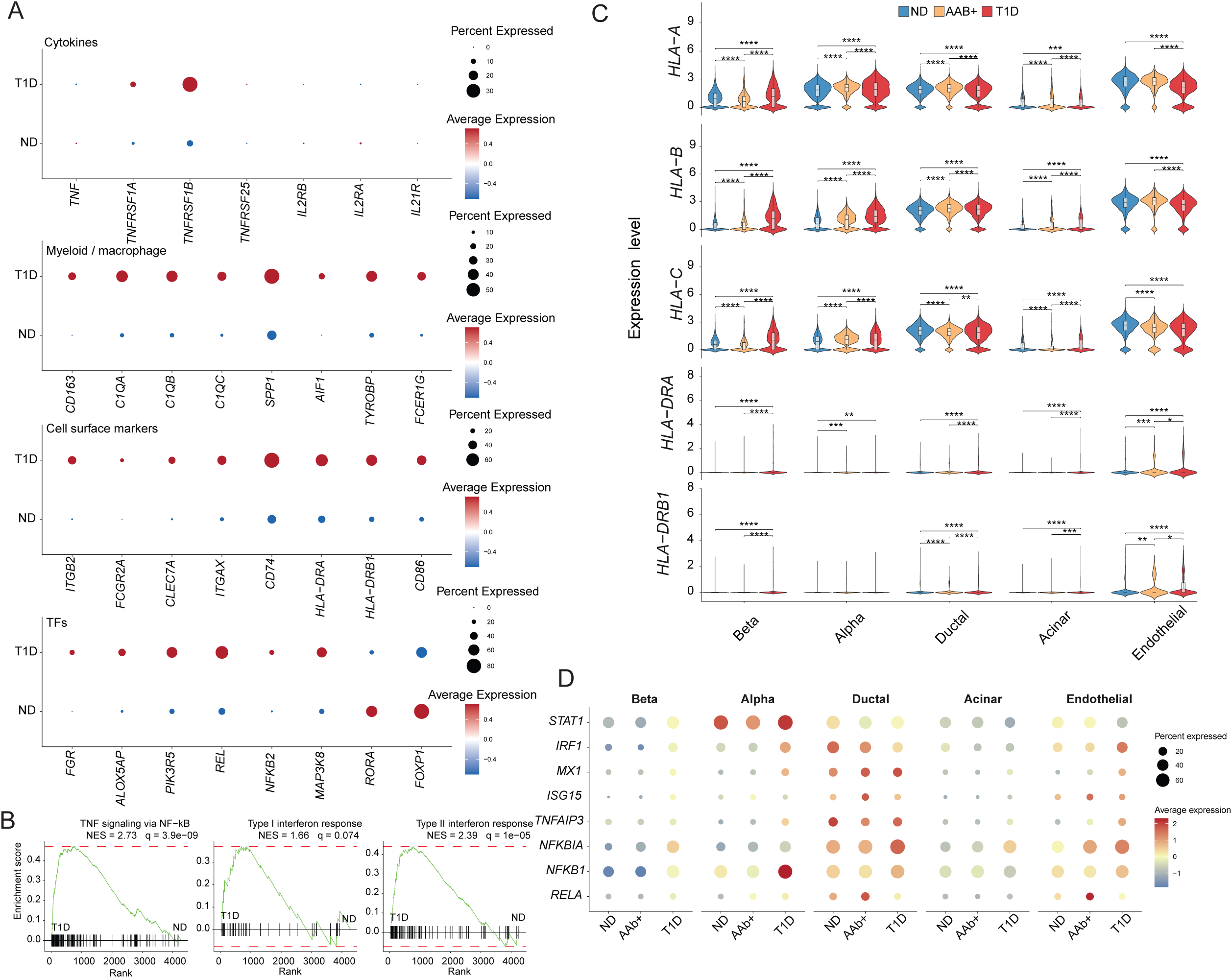
T1D-associated immune remodeling and inflammatory responses across pancreatic cell populations. (A) Dot plots comparing expression of selected cytokine, myeloid/macrophage, cell-surface, and transcriptional-regulator genes between ND and T1D immune cells. Dot size represents the percentage of cells expressing each gene, and color represents scaled average expression. (B) Gene set enrichment analysis (GSEA) of genes ranked by differential expression between T1D and ND immune cells. Enrichment plots show TNF signaling via NF-κB (NES = 2.73, *q* = 3.9 × 10⁻⁹), type I interferon response (NES = 1.66, *q* = 0.074), and type II interferon response (NES = 2.39, *q* = 1.0 × 10⁻⁵). GSEA significance is reported using FDR-adjusted *q* values. (C) Violin plots showing normalized single-cell RNA expression of *HLA-A, HLA-B, HLA-C, HLA-DRA,* and *HLA-DRB1* across β, α, ductal, acinar, and endothelial cells from ND, AAb+, and T1D samples. Internal boxplots indicate the median and interquartile range. Pairwise differential expression between ND and AAb+, ND and T1D, and AAb+ and T1D was assessed separately within each cell type using MAST, with cellular gene detection rate included as a covariate. *P* values were adjusted across gene, cell-type, and group comparisons using the Benjamini–Hochberg false discovery rate (FDR) procedure. Brackets indicate statistically significant comparisons (FDR ≤ 0.05); non-significant comparisons are not shown. Significance is denoted as FDR ≤ 0.05 (*), ≤ 0.01 (**), ≤ 0.001 (***), and ≤ 0.0001 (****). (D) Dot plots showing expression of selected interferon-associated (*STAT1, IRF1, MX1,* and *ISG15*) and TNF/NF-κB-associated (*TNFAIP3, NFKBIA, NFKB1,* and *RELA*) genes across β, α, ductal, acinar, and endothelial cells from ND, AAb+, and T1D samples. Dot size represents the percentage of cells with detectable expression of each gene, and color represents scaled average SCT-normalized expression within each gene.

We next examined genes involved in immune-cell surface signaling and antigen presentation. T1D immune cells showed increased or altered expression of *ITGB2, FCGR2A, CLEC7A, ITGAX, CD74, HLA-DRA, HLA-DRB1,* and *CD86* (Fig. 3A). Increased expression of *CD74* and HLA class II-associated genes is consistent with enhanced antigen presentation, whereas the *FCGR2A, CLEC7A,* and *ITGAX* pattern suggests a greater contribution of myeloid cells. We also observed differences in TFs and intracellular signaling regulators. T1D immune cells showed increased expression of inflammatory signaling genes, including *FGR, ALOX5AP, PIK3R5, REL, NFKB2,* and *MAP3K8*, whereas regulatory factors such as *RORA* and *FOXP1* were more prominent in ND cells (Fig. 3A). *REL* and *NFKB2*, both central components of NF-κB signaling, were among the T1D-enriched regulators, and the concurrent enrichment of *FGR, ALOX5AP,* and *PIK3R5* further supports activation of myeloid-associated signaling pathways. These data point to a coordinated shift in T1D immune cells involving cytokine signaling, macrophage-associated genes, antigen presentation, and NF-κB-related transcriptional regulation.

To determine whether these transcriptional alterations converged on specific inflammatory pathways, we performed gene set enrichment analysis (GSEA) using genes ranked by differential expression between T1D and ND immune cells. TNF signaling through NF-κB was strongly enriched toward the T1D-associated end of the ranked gene list (NES = 2.73, *q* = 3.9 × 10⁻⁹; Fig. 3B). The type II interferon response was also significantly enriched toward T1D (NES = 2.39, *q* = 1.0 × 10⁻⁵), whereas the type I interferon response showed a weaker positive enrichment that did not reach the FDR threshold of *q* < 0.05 (NES = 1.66, *q* = 0.074). These results indicate that the immune-cell transcriptional changes in T1D are accompanied by coordinated activation of TNF–NF-κB and type II interferon programs.

We next asked whether these disease-associated inflammatory changes extended beyond the immune compartment. Given the established role of altered antigen presentation in T1D, and previous reports of HLA class I hyperexpression in pancreatic islets from individuals with T1D and from autoantibody-positive donors^25–27^, we examined HLA expression across major pancreatic cell populations. We compared single-cell RNA expression of the HLA class I genes *HLA-A, HLA-B,* and *HLA-C* and the HLA class II genes *HLA-DRA* and *HLA-DRB1* across β, α, ductal, acinar, and endothelial cells from ND, AAb+, and T1D samples (Fig. 3C). HLA class I genes were broadly expressed across pancreatic cell populations and showed significant disease-associated differences in multiple cell types by MAST^28^ after multiple-testing correction (BH-adjusted *P* ≤ 0.05; Fig. 3C), with many pairwise comparisons reaching adjusted *P* ≤ 0.0001. The magnitude and direction of these differences varied across genes and cell types, indicating a heterogeneous rather than uniform HLA class I transcriptional response.

In contrast, *HLA-DRA* and *HLA-DRB1* showed a more restricted and heterogeneous expression pattern. These transcripts were sparse in many β, α, and acinar cells and more evident in subsets of cells, particularly in ductal and endothelial populations (Fig. 3C). Despite this overall sparsity, significant disease-associated differences in *HLA-DRA* and/or *HLA-DRB1* expression were detected in several pancreatic cell populations by MAST^28^ (BH-adjusted *P* ≤ 0.05), with several comparisons reaching adjusted *P* ≤ 0.001 or ≤ 0.0001 (Fig. 3C). These findings indicate broad disease-associated modulation of HLA class I transcription across pancreatic cell types, whereas HLA class II expression was more restricted and heterogeneous.

Together, these analyses reveal coordinated but heterogeneous inflammatory remodeling in the T1D islets. A T1D-associated myeloid/macrophage transcriptional phenotype coincided with strong TNF–NF-κB and type II interferon pathway enrichment in immune cells and with altered HLA-ABC and HLA-DR expression across multiple non-immune pancreatic populations. These findings support a model in which T1D-associated immune remodeling is accompanied by broader inflammatory transcriptional changes across endocrine, exocrine, and vascular compartments of the pancreatic microenvironment.

Given the enrichment of TNF–NF-κB and interferon signaling in T1D immune cells, we next examined whether components of these inflammatory programs were also altered across non-immune pancreatic cell populations. Expression of selected interferon-associated genes (*STAT1, IRF1, MX1,* and *ISG15*) and TNF/NF-κB-associated genes (*TNFAIP3, NFKBIA, NFKB1,* and *RELA*) was evaluated across β, α, ductal, acinar, and endothelial cells from ND, AAb+, and T1D samples (Fig. 3D). These genes displayed marked cell type-and disease-dependent expression patterns. In particular, α cells showed increased expression of several inflammatory signaling genes in T1D, including *STAT1* and *NFKB1*, whereas ductal cells exhibited prominent expression of *IRF1, MX1, TNFAIP3,* and *NFKBIA*. Endothelial cells also showed disease-associated changes in inflammatory-response genes, including *IRF1, ISG15, NFKBIA,* and *RELA*. By contrast, β and acinar cells generally displayed more restricted or heterogeneous responses.

The inflammatory transcriptional patterns were not uniform across disease stages or cell types. AAb+ cells showed intermediate or distinct expression patterns for several genes, including *NFKBIA* and *RELA* in endothelial cells and selected interferon-associated genes in ductal and acinar populations. These observations suggest that inflammatory signaling outside the immune compartment is heterogeneous and may involve different transcriptional programs in different pancreatic cell populations rather than a uniform tissue-wide response.

Together with the HLA-ABC and HLA-DR alterations observed in Fig. 3C, these findings indicate that the inflammatory phenotype associated with T1D extends beyond infiltrating immune cells. The concordance between TNF–NF-κB/interferon pathway enrichment in immune cells and altered expression of downstream inflammatory-response genes in endocrine, exocrine, and endothelial populations supports broader remodeling of the pancreatic inflammatory microenvironment.

### ChromBPNet predicts cell type-specific regulatory effects of T1D-associated variants

To examine individual variants with potential regulatory effects, we next asked whether T1D-associated genetic variants alter chromatin accessibility in a cell type-specific manner. We obtained T1D GWAS data from the NHGRI-EBI GWAS Catalog^29^; the search returned 101 GWAS studies of type 1 diabetes mellitus (see Methods). We expanded lead T1D GWAS variants to variants in linkage disequilibrium (LD; *r*² > 0.8) and intersected these variants with cell type-specific accessible chromatin regions identified by scATAC-seq (Fig. 4A). We then applied cell type-specific ChromBPNet^19^ models to predict the sequence-dependent effect of each candidate variant on local chromatin accessibility. This approach goes beyond overlap between GWAS variants and accessible regions and estimates the regulatory consequences of alternative alleles within the chromatin landscape of individual pancreatic cell types (Fig. 4A).

**Figure 4.**
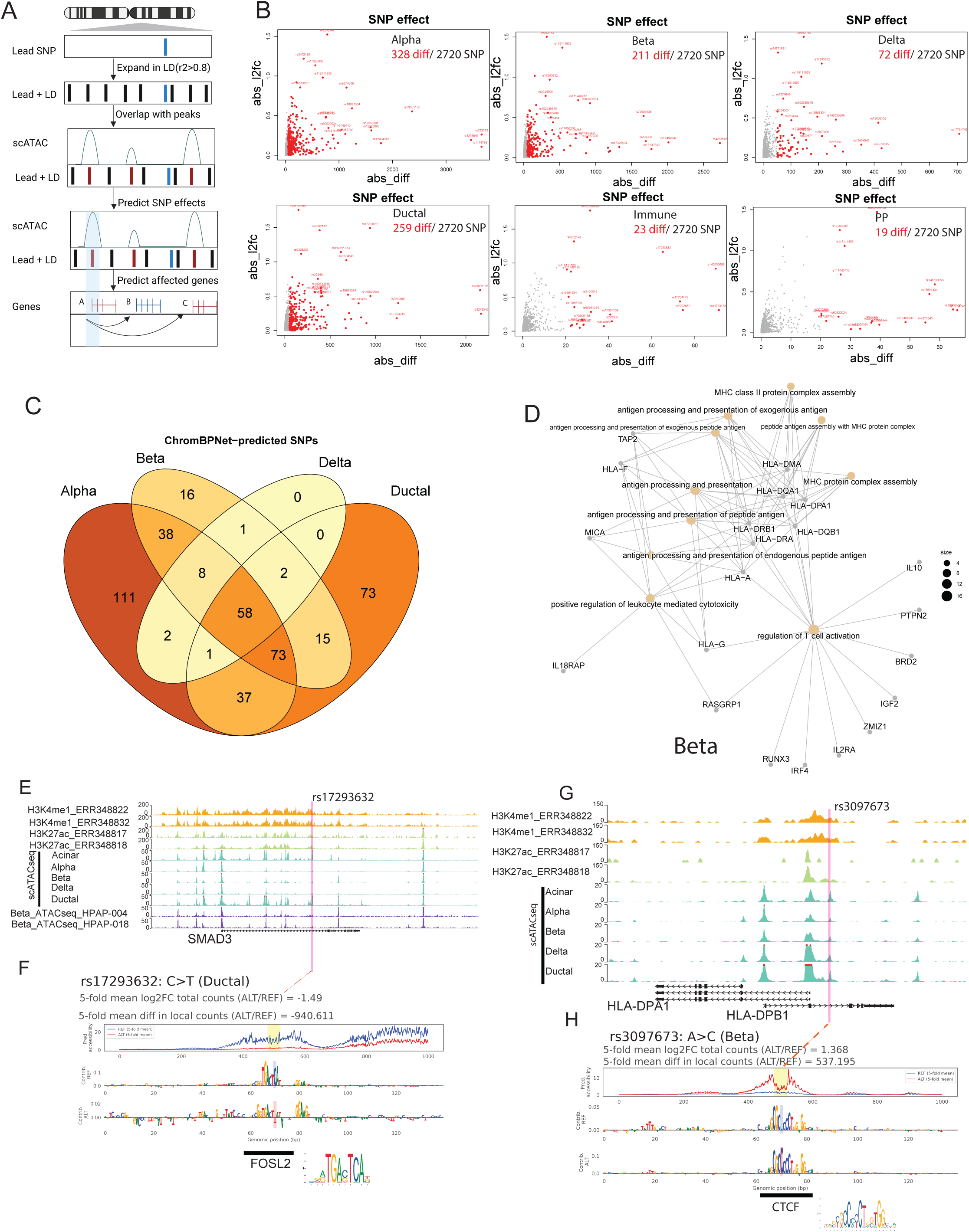
ChromBPNet identifies T1D-associated variants with predicted cell type-specific effects on chromatin accessibility. (A) Schematic of the integrative workflow used to prioritize candidate T1D-associated regulatory SNPs and link them to potential target genes. T1D-associated variants overlapping cell type-specific accessible chromatin regions were evaluated using ChromBPNet to predict allele-dependent effects on chromatin accessibility and to prioritize candidate regulatory variants. (B) Number of T1D-associated SNPs in each pancreatic islet cell type with large ChromBPNet-predicted effects on chromatin accessibility, defined as an absolute mean difference in local counts (ALT/REF) > 50 across the five cross-validation folds. (C) Overlap of predicted regulatory SNPs across α, β, δ, and ductal cells, highlighting shared and cell type-specific variant effects. (D) Gene Ontology (GO) enrichment analysis of genes associated with SNPs predicted to alter β-cell chromatin accessibility. Representative SNP-associated genes contributing to enriched biological processes are indicated. (E–F) Regulatory characterization of rs17293632. The variant overlaps a regulatory region marked by H3K4me1 and H3K27ac and is located within accessible chromatin, with particularly strong accessibility in ductal cells (E). ChromBPNet predicts an allele-dependent effect of rs17293632 on local chromatin accessibility, and sequence-based motif analysis implicates FOSL2 as a candidate transcription factor associated with the variant-containing sequence (F). (G–H) Regulatory characterization of rs3097673 near HLA-DPB1. The variant is located within a β-cell-accessible regulatory region (G), and ChromBPNet predicts that the alternative allele increases local chromatin accessibility in β cells. Sequence-based motif analysis implicates CTCF as a candidate transcription factor associated with the variant-containing sequence (H).

We first quantified the predicted allelic effects of candidate variants across pancreatic cell populations. ChromBPNet identified subsets of the tested variants with predicted effects on chromatin accessibility in each cell type, but the number and magnitude of these effects differed substantially across cell types (Fig. 4B). Among the 2,720 variants evaluated, 328 variants in α cells, 211 in β cells, 72 in δ cells, 259 in ductal cells, 23 in immune cells, and 19 in PP cells had an absolute mean difference in local counts (ALT/REF) > 50 across the five cross-validation folds, a threshold used to highlight variants with larger predicted accessibility effects (red points in Fig. 4B). These results indicate that T1D-associated regulatory variation is not confined to β cells and instead includes substantial predicted effects in α and ductal cells, together with smaller sets of variants with predicted effects in other pancreatic populations.

We next asked whether the same variants were predicted to alter accessibility across multiple endocrine and non-endocrine cell types or whether their effects were predominantly cell type-specific. Comparison of ChromBPNet-prioritized variants across α, β, δ, and ductal cells revealed both shared and cell type-restricted regulatory effects (Fig. 4C). A subset of variants was shared across several populations, including 58 variants common to all four cell types, whereas substantial numbers were restricted to individual populations, including 111 α-cell-specific and 73 ductal-cell-specific variants. Other variants were shared between selected combinations of cell types. Thus, although every cell carries the same genetic variants, their predicted effects on chromatin accessibility can depend strongly on the underlying cell type-specific regulatory landscape.

To investigate the potential biological consequences of these regulatory variants, we linked β-cell candidate regulatory elements to putative target genes and examined the functional relationships among these genes (Fig. 4D). The resulting network highlighted genes associated with antigen processing and presentation and HLA class II-related functions, including *HLA-DPA1, HLA-DPB1, HLA-DRA,* and *HLA-DRB1*, together with additional genes involved in immune and cellular regulatory processes. This analysis suggests that a subset of T1D-associated variants acting within β-cell accessible chromatin may influence antigen-presentation-associated processes.

We then examined individual variants to determine how sequence variation could generate these cell type-specific regulatory effects. The T1D-associated variant rs17293632 overlapped an active regulatory region marked by H3K4me1 and H3K27ac that was accessible across pancreatic cell populations, with prominent accessibility in ductal cells (Fig. 4E). ChromBPNet predicted a strong allele-dependent effect of rs17293632 in ductal cells, with the C>T substitution predicted to reduce accessibility (mean across five folds: log2 fold change = −1.49, differential contribution score = −940.611; Fig. 4F). Nucleotide-level contribution scores localized the predicted regulatory effect to the variant-containing sequence, and motif analysis implicated FOSL2, suggesting that altered binding of AP-1-family TFs may contribute to the accessibility effect at this locus.

rs3097673, located within the HLA class II region near *HLA-DPA1* and *HLA-DPB1*, points to a different predicted regulatory mechanism (Fig. 4G). In contrast to rs17293632, ChromBPNet predicted that the alternative allele of rs3097673 increases local chromatin accessibility in β cells (mean across five folds: log2 fold change = 1.368, differential contribution score = 537.195; Fig. 4H). Nucleotide-resolution contribution scores showed a strong sequence-dependent effect around the variant, and motif analysis implicated CTCF at the affected sequence. Together with the β-cell target-gene network enriched for HLA class II and antigen-presentation genes, this locus provides an example of how T1D-associated genetic variation may influence β-cell regulatory activity within an immune-relevant genomic region.

Overall, these analyses indicate that T1D-associated variants can have strongly context-dependent predicted effects on pancreatic chromatin accessibility. ChromBPNet identified both shared and cell type-restricted regulatory variants, with the largest numbers of predicted effects in α, ductal, and β cells. Base-resolution analysis of rs17293632 and rs3097673 further illustrates how individual risk variants may perturb distinct sequence-specific regulatory mechanisms involving FOSL2/AP-1 and CTCF, respectively. These findings extend the cell type-level GWAS analysis by showing that, even in the absence of significant heritability enrichment within a single pancreatic cell type, individual T1D-associated variants can have pronounced and highly cell type-specific predicted regulatory effects.

### Integration of ChromBPNet and transcription factor footprinting identifies candidate variant–TF regulatory mechanisms

Having identified T1D-associated variants with predicted effects on chromatin accessibility, we next asked whether these variants could perturb sequence-specific TF activity. We therefore integrated cell type-specific ChromBPNet predictions with TF footprinting using scPrinter^30^. ChromBPNet provided sequence-based estimates of allele-dependent changes in accessibility, whereas multiscale footprinting of bulk ATAC-seq data from β cells of nine PANC-DB donors (Supplementary Table 2) enabled us to examine local Tn5 insertion patterns and nominate TFs with predicted binding activity near variant-containing regulatory elements (Fig. 5). Because the footprinting data served as a β-cell regulatory reference rather than as a measure of disease-specific TF occupancy, we interpreted these analyses as evidence of local TF-associated regulatory activity. This combined analysis allowed us to prioritize candidate variant–TF relationships supported by both sequence-dependent accessibility effects and local chromatin footprinting.

**Figure 5.**
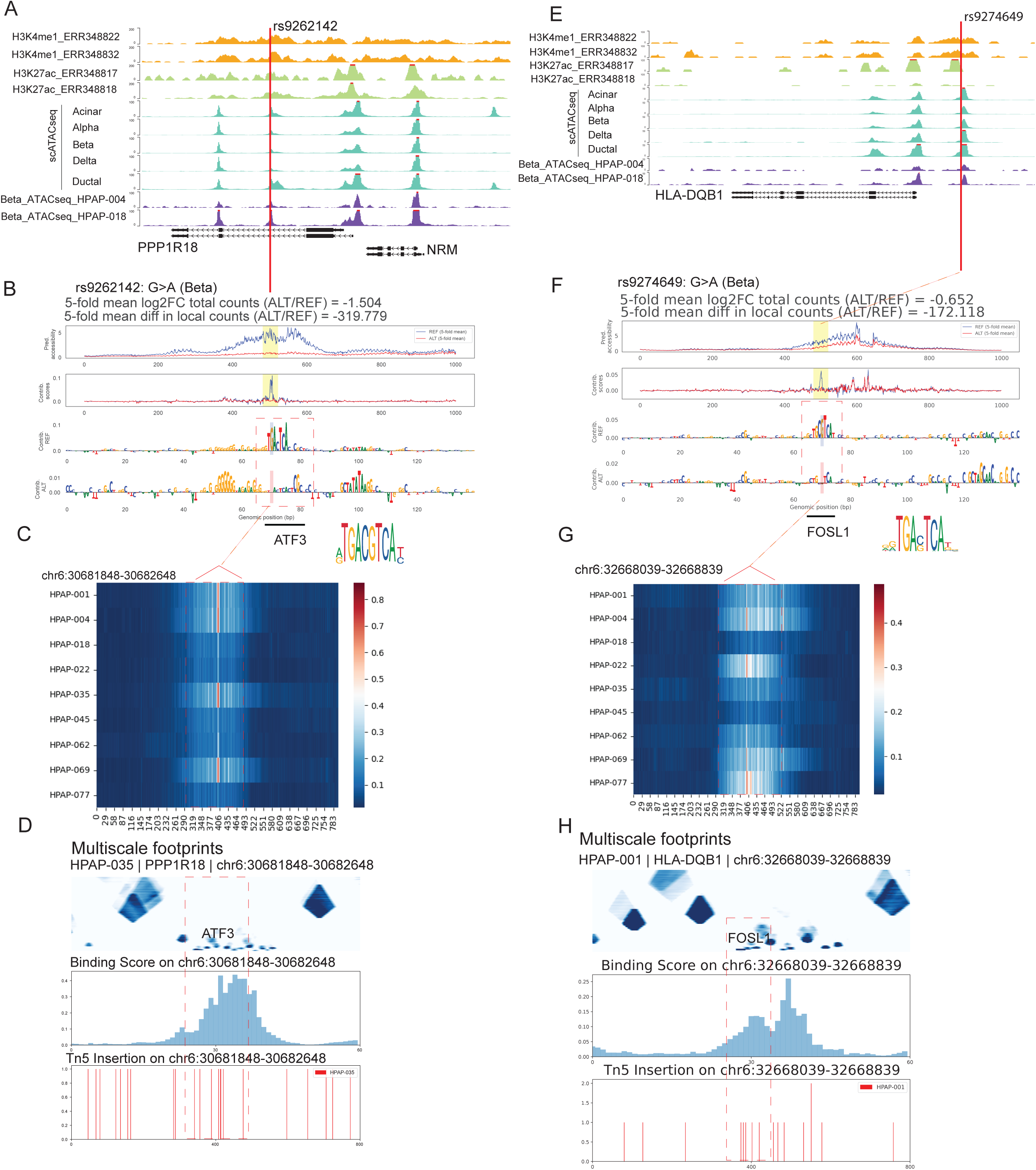
Integration of ChromBPNet and transcription factor footprinting identifies candidate regulatory mechanisms at T1D-associated β-cell variants. (A–D) Regulatory characterization of rs9262142 in β cells. (A) Chromatin landscape surrounding rs9262142, showing its localization within a β-cell-accessible regulatory region. (B) ChromBPNet-predicted allele-dependent effects of rs9262142 on local chromatin accessibility. (C) scPrinter footprinting analysis across the rs9262142-containing regulatory region in β-cell ATAC-seq data. (D) Multiscale footprinting, transcription factor (TF) binding scores, and Tn5 insertion profiles surrounding rs9262142, identifying ATF3 as a candidate TF with a footprint near the variant. (E–H) Regulatory characterization of rs9274649 near HLA-DQB1 in β cells. (E) Chromatin landscape surrounding rs9274649, showing its localization within a β-cell-accessible regulatory region. (F) ChromBPNet-predicted allele-dependent effects of rs9274649 on local chromatin accessibility. (G) scPrinter footprinting analysis across the rs9274649-containing regulatory region in β-cell ATAC-seq data. (H) Multiscale footprinting, TF binding scores, and Tn5 insertion profiles surrounding rs9274649, identifying FOSL1 as a candidate TF with a footprint near the variant.

We first examined rs9262142, which overlapped an accessible regulatory region in β cells and showed an allele-dependent effect on local chromatin accessibility predicted by ChromBPNet (Fig. 5A,B). Multiscale footprinting revealed a localized footprint around the variant-containing region, indicating that rs9262142 lies within a region whose accessibility pattern is compatible with sequence-specific TF binding (Fig. 5C). We next examined TF binding scores and local Tn5 insertion profiles to identify candidate factors associated with this regulatory element. ATF3 showed a footprint near rs9262142, nominating ATF3 as a candidate TF whose activity may be sensitive to sequence variation at this locus (Fig. 5C,D). Together, the sequence-based accessibility prediction and footprinting results suggest that rs9262142 may alter local regulatory activity within an ATF3-associated regulatory element.

We next examined rs9274649, located near the T1D-associated *HLA-DQB1* locus. Like rs9262142, rs9274649 overlapped a β-cell-accessible regulatory region and was predicted by ChromBPNet to produce an allele-dependent change in local chromatin accessibility (Fig. 5E,F). scPrinter analysis identified a localized footprint and TF binding signal around the variant-containing sequence (Fig. 5G,H). Among the candidate factors associated with this region, FOSL1 showed a footprint proximal to rs9274649, nominating a potential rs9274649–FOSL1 regulatory interaction. FOSL1 is an AP-1 family TF, which links this candidate variant to an AP-1-associated regulatory program and to the AP-1 motif enrichment observed independently in cCREs that gained accessibility in disease (Fig. 2E).

The identification of an AP-1-associated factor at rs9274649 is consistent with our broader multiomic analyses. T1D-associated gained cCREs showed enrichment of AP-1 and NF-κB family motifs (Fig. 2E), and transcriptional analyses identified enrichment of TNF–NF-κB and inflammatory-response programs. The FOSL1-associated footprint at rs9274649 therefore provides a locus-specific example in which a genetically associated variant, cell type-specific accessible chromatin, sequence-predicted regulatory activity, and an inflammatory TF-associated footprint converge within the same regulatory region.

## Discussion

In this study, we integrated single-cell multiomic profiling, T1D genetic association data, sequence-based deep learning^19^, and transcription factor (TF) footprinting^30^ to investigate regulatory alterations in human pancreatic islets and to prioritize noncoding variants with potential regulatory effects. We identified widespread but heterogeneous changes in gene expression and chromatin accessibility across pancreatic cell populations in T1D. By combining these disease-associated regulatory landscapes with cell type-specific sequence modeling, we further identified candidate T1D-associated variants predicted to alter chromatin accessibility and nominated TFs that may contribute to their regulatory effects. These analyses provide a framework for examining T1D-associated noncoding variation within the cellular and regulatory contexts in which it may function.

A prominent feature of the T1D islet landscape was the upregulation of inflammatory and stress-associated transcriptional programs. Interferon-response pathways were among the programs increased in T1D, whereas pathways related to secretory, metabolic, and cell identity-associated functions were reduced. Chromatin accessibility showed extensive remodeling in parallel, with both gains and losses of cCREs across pancreatic cell populations. Regulatory elements gaining accessibility were enriched for TF motifs associated with inflammatory and stress-responsive programs, including AP-1 and NF-κB, whereas elements losing accessibility showed a distinct repertoire of regulatory motifs, including RFX-family motifs. The correspondence between these transcriptional and chromatin signatures suggests that the altered transcriptional state of T1D islets occurs in the context of broad reorganization of the underlying regulatory landscape.

The magnitude of these changes differed among pancreatic cell populations. Although β cells are central to T1D pathogenesis, transcriptional and chromatin alterations were also observed across other endocrine and non-endocrine populations. This broader response is consistent with the view of the pancreatic islet as a multicellular tissue in which autoimmune and inflammatory stress can influence multiple cellular compartments^6,21,31,32^. At the same time, differences in the extent and nature of regulatory remodeling among cell populations emphasize the importance of resolving disease-associated changes at the cell-type level rather than relying exclusively on whole-islet measurements.

Cell type-specific ChromBPNet models provided a complementary approach by predicting how allelic sequence changes may alter local chromatin accessibility^19^. Several candidate variants showed predicted effects that differed across cell types, illustrating how the regulatory consequence of the same sequence variant may depend on the surrounding cellular regulatory environment. For example, rs17293632 localized to an active regulatory region and showed a predicted allele-dependent accessibility effect in ductal cells, with motif analysis implicating FOSL2 as a candidate regulator. In contrast, rs3097673, located near *HLA-DPB1*, showed a predicted effect on chromatin accessibility in β cells and was associated with a CTCF motif. These examples point to distinct candidate mechanisms through which T1D-associated noncoding variation may influence regulatory activity in endocrine and non-endocrine pancreatic cells.

Integrating sequence-based predictions with TF footprinting provided an additional layer for prioritizing potential variant–TF relationships. At rs9262142, the predicted allele-dependent effect on β-cell chromatin accessibility coincided with a localized footprint implicating ATF3, whereas rs9274649 near *HLA-DQB1* was associated with a predicted accessibility effect and a footprint implicating FOSL1. FOSL1 and FOSL2 both belong to the AP-1 family, linking two variant-level observations to the broader stress-and inflammation-associated regulatory signatures detected in T1D islets. This convergence raises the possibility that AP-1-related regulatory activity contributes to T1D pathogenesis in multiple pancreatic cell types. Our study has several limitations. ChromBPNet predicts sequence-dependent accessibility, and footprinting infers potential TF occupancy; neither approach directly demonstrates allele-specific TF binding. These variant–TF relationships should therefore be considered candidate mechanisms that require experimental validation. In addition, the study relies primarily on publicly available human pancreatic datasets generated from different donors and experimental settings, and the limited number of T1D and autoantibody-positive samples restricts statistical power after stratification by cell type. Inter-individual variation, disease duration, and technical differences may also contribute to the observed regulatory heterogeneity.

In summary, our study defines cell type-resolved transcriptional and chromatin remodeling in human pancreatic islets in T1D and places disease-associated noncoding variation within these regulatory landscapes. Integration with TF motif and footprinting analyses further nominated specific variant–TF relationships, including AP-1-associated regulatory mechanisms, for functional investigation. These findings provide a path from genetic association to experimentally testable regulatory hypotheses and highlight the importance of considering both endocrine and non-endocrine pancreatic cell populations when investigating the regulatory consequences of T1D-associated variation.

## Methods

### Integration of Multiomic Data Across HPAP Samples

Single-cell multiome (scRNA-seq and scATAC-seq) data were downloaded from the PANC-DB database (https://hpap.pmacs.upenn.edu). Metadata on the generation and sequencing of each sample are available on the PANC-DB website via the ‘Download All Metadata’ button on the ‘Experiment Data Download’ page. We selected 20 high-quality samples from ND, AAb+, and T1D donors (Supplementary Table 1). Cell Ranger ARC (v2.0.2) was used to demultiplex the sequencing output and generate two cell-by-feature matrices per sample, with reads aligned to the GRCh38-2024-A reference genome provided by 10x Genomics. Quality control was performed separately for each modality. Low-quality cells were excluded using the following thresholds: fewer than 200 or more than 25,000 total RNA counts, fewer than 1,000 or more than 70,000 peak counts, or a mitochondrial read fraction above 20%. Doublets were detected and removed using DoubletFinder^33^. For cell type annotation, we integrated cells across samples using Seurat^34^ and Signac (v1.17.0)^35^, embedding all cells into a shared low-dimensional space based on both gene expression and chromatin accessibility profiles. RNA counts were normalized with SCTransform, with the per-cell mitochondrial percentage regressed out. Principal component analysis (PCA) was performed on the RNA data, and UMAP was run on the first 50 principal components (PCs); the number of PCs (50) was chosen using an elbow plot. scATAC-seq peak counts were normalized with term frequency–inverse document frequency (TF-IDF), and dimension reduction was performed by singular value decomposition (SVD) of the normalized peak matrix (latent semantic indexing, LSI). The scATAC-seq UMAP was computed from LSI components 2 through 50. A weighted nearest neighbor (WNN) graph representing a weighted combination of both modalities was constructed with Seurat’s FindMultiModalNeighbors, using the first 50 dimensions of the scRNA-seq reduction and dimensions 2 through 50 of the scATAC-seq reduction.

BigWig files were generated from merged BAM files using the bamCoverage utility in deepTools^36^ with the parameters -e 100 -bs 100 --normalizeUsing RPKM. Density plots were produced with the Nebulosa^37^ package.

### HLA gene expression analysis

Expression of *HLA-A, HLA-B, HLA-C, HLA-DRA,* and *HLA-DRB1* was compared across ND, AAb+, and T1D groups in β, α, ductal, acinar, and endothelial cells. Differential expression was assessed separately within each cell type using MAST^28^ as implemented in Seurat FindMarkers, with cellular gene detection rate included as a covariate. Pairwise comparisons were performed between ND and AAb+, ND and T1D, and AAb+ and T1D. *P* values were adjusted for multiple testing using the Benjamini– Hochberg false discovery rate (FDR) procedure, with FDR ≤ 0.05 considered statistically significant.

### Gene set enrichment analysis

Gene set enrichment analysis (GSEA) was performed using the fgsea^38^ R package. Genes from the T1D-versus-ND immune-cell differential-expression analysis were ranked according to average log2 fold change, with positive values corresponding to higher expression in T1D and negative values corresponding to higher expression in ND. Pathway enrichment was summarized using the normalized enrichment score (NES), nominal *P* value, and false-discovery rate (FDR)-adjusted *q* value. TNF signaling via NF-κB, interferon-α response, and interferon-γ response pathways were selected for visualization based on their relevance to the inflammatory transcriptional changes observed in the immune compartment. For GSEA, FDR-adjusted *q* values < 0.05 were considered statistically significant.

### ChIP-seq analysis

ChIP-seq data were downloaded from the Sequence Read Archive (SRA), including H3K4me1 datasets (ERR348822, ERR348832) and H3K27ac datasets (ERR348817, ERR348818). Bowtie2 (v2.3.3.1; --very-sensitive mode) was used to map ChIP-seq reads to the human reference genome GRCh38. Duplicate reads were removed using the MarkDuplicates function from Picard tools v.2.17.0 (http://broadinstitute.github.io/picard/). Reads per kilobase per million mapped reads (RPKM)-normalized bigWig files were generated with bamCoverage from deepTools (v3.3.2)^36^.

### GWAS analysis

T1D GWAS data were obtained from the NHGRI-EBI GWAS Catalog^29^. The search returned 101 GWAS studies of type 1 diabetes mellitus. Of the 1212 associations in the GWAS Catalog, 1007 met the *P*-value threshold of 1×10^−6^, corresponding to 421 genes. For each locus, we used the lead variant and identified all SNPs in high LD with it (R^2^ > 0.8) using the LDproxy tool from the LDlinkR package (v1.4.0)^39^.

### ATAC–seq data processing

Bulk ATAC-seq data from β cells of nine donors were downloaded from the PANC-DB database (https://hpap.pmacs.upenn.edu) (Supplementary Table 2). Read quality was assessed using FastQC (version 0.11.8) (http://www.bioinformatics.babraham.ac.uk/projects/fastqc/). Adapters were removed using Trim Galore (v0.6.6), and trimming quality was verified using FastQC v0.11.8. Reads were aligned to the GRCh38 reference genome assembly using Bowtie2 (v2.5.0), and alignments were saved as BAM files. Duplicate reads were identified using Picard MarkDuplicates. Genome browser tracks were generated with deepTools (v3.3.2)^36^ bamCoverage using the --normalizeUsing RPKM option and visualized in the WashU Epigenome Browser (https://epigenomegateway.wustl.edu/). Peak calling (*q* < 0.05) was performed with MACS2 (v2.2.9.1)^40^, using the BAMPE model and removing all duplicate reads.

### ChromBPNet model training and deriving contribution scores

We analyzed scATAC-seq data using ChromBPNet. To define genomic regions for training ChromBPNet models^19^, we performed peak calling to obtain a lenient set of accessible regions. First, pseudobulk fragment files were generated for each cell type cluster by concatenating scATAC-seq fragments from all cells in that cluster across all samples. MACS2 (v2.2.9.1) was used to call peaks on the pseudobulk files with the parameters -g hs --nomodel --keep-dup all --call-summits. Peaks overlapping the GRCh38 ENCODE blacklist (ENCODE accession ENCFF356LFX) were excluded. Peak coordinates were resized to 1,000 bp centered on the MACS2 peak summit. We used the ChromBPNet package (https://github.com/kundajelab/chrombpnet, v1.0.1) and followed the workflow described in its documentation. We used the command chrombpnet prep nonpeaks to define background regions matched to the GC content of peak regions. ChromBPNet takes as input one-hot-encoded 2114 bp DNA sequences centered on ATAC-seq peaks. The model is trained to predict two outputs: the total Tn5 insertion counts over a 1000 bp central region and the base-resolution Tn5 insertion profile across this region. For each cell type, we used a fivefold cross-validation scheme in which each fold (designated 0 to 4) comprised a different set of training, validation, and test chromosomes, with each chromosome included in the test set of at least one fold. The test and validation chromosomes for each fold were as follows: fold 0 (test: chr1, chr3, chr6; validation: chr8, chr19), fold 1 (test: chr2, chr8, chr17; validation: chr5, chr10, chr18), fold 2 (test: chr5, chr9, chr11; validation: chr12, chr14), fold 3 (test: chr4, chr10, chr18; validation: chr2, chr7), and fold 4 (test: chr5, chr15, chr12; validation: chr3, chr6).

To interpret model predictions, we applied the DeepLIFT algorithm^41^, which quantifies the contribution of each base to the predicted chromatin accessibility. Contribution scores were calculated as the additive decomposition of the difference between the predicted log(total counts) of the native sequence and the average prediction across 20 dinucleotide-shuffled background sequences. Scores were computed for each of the five fold-specific models and averaged, and the averaged scores were used for de novo TF motif discovery. Motif discovery was conducted using TF-MoDISco^42^ (https://github.com/kundajelab/tfmodisco), which clusters high-contribution subsequences (seqlets) to identify representative, nonredundant motifs across accessible chromatin regions. Motif discovery was performed over 500-bp windows centered on peak summits, with a maximum of 50,000 seqlets considered per metacluster. Identified motifs were annotated to putative TFs or TF families by comparison with motifs from the MEME v4 database using TOMTOM^43^, with the command tomtom -no-ssc -oc . --verbosity 1 -text -min-overlap 5 -mi 1 -dist pearson -evalue -thresh 10.0. Finally, to map predictive instances of the de novo motifs, we scanned contribution score tracks using FiNeMo (https://github.com/kundajelab/Fi-NeMo). We ran FiNeMo with the complete set of de novo contribution weight matrices (CWMs) discovered by TF-MoDISco, setting --global-lambda 0.7 and --cwm-trim-threshold 0.3 and using default values for the remaining parameters. These high-confidence motif instances were later intersected with memory domains to evaluate their role in establishing long-term accessibility.

### Prediction of variant effects using ChromBPNet models

We used the tangermeme package (https://github.com/jmschrei/tangermeme) for predictions and model interpretation. For each variant, we used the 1,000 bp model training peaks for the relevant cell type to extract the reference genome sequence of the peak that the variant overlapped. This sequence (extended equally on either side to 2,114 bp) was input to all five fold-specific ChromBPNet models for the relevant cell type to obtain the predicted accessibility profile and aggregate log counts in the peak. For each fold, predicted profile logits were converted into accessibility profiles by applying a softmax and scaling by the exponentiated predicted log counts, and model interpretation with respect to the counts output was performed using DeepLIFT. Next, the effect allele was substituted into the sequence at the variant position, and predictions and contribution scores were obtained as for the reference sequence. For each model, we computed the variant effect as the sum of differences in per-base predicted read counts in the 100 bp window centered on the variant, and we then averaged the effect score across folds. We also computed the log2 fold change between predicted counts for the effect versus the non-effect allele over the peak region, where a log2 fold change > 0 indicates that the effect allele was predicted to increase accessibility. Figures show the mean predicted profiles and contribution scores across folds.

### Footprinting and seq2PRINT

Bulk ATAC-seq datasets from β cells (Supplementary Table 2) were used to train the scPrinter models. Footprint scores were calculated using the default PRINT v3 procedure described at https://ruochiz.com/scprinter_doc/index.html (scPrinter, the Python implementation of PRINT^30^). Multiscale footprint scores were computed using the scp.tl.get_footprint_score function across spatial scales of 2-100 bp with parallel processing (n_jobs = 32), and the resulting footprint score matrices were stored in backed mode. These matrices were then used to visualize multiscale TF footprints across genomic regions and cell groups with the scp.pl.plot_footprints function.

### Genome browser visualizations

All genomic tracks are hosted online for interactive visualization with the WashU Genome Browser (https://epigenomegateway.wustl.edu/browser2022/?genome=hg38%20&hub=https://human-dev-multiome-atlas.s3.amazonaws.com/tracks/%20HDMA_trackhub.json).

## Data Availability

ChIP-seq data were downloaded from the Sequence Read Archive (SRA), including H3K4me1 datasets (ERR348822, ERR348832) and H3K27ac datasets (ERR348817, ERR348818). Bulk ATAC-seq datasets from β cells (HPAP-001, HPAP-004, HPAP-018, HPAP-022, HPAP-035, HPAP-045, HPAP-062, HPAP-069, and HPAP-077), as well as single-cell multiomic datasets (scATAC-seq and scRNA-seq; HPAP-093, HPAP-097, HPAP-104, HPAP-130, HPAP-137, HPAP-139, HPAP-141, HPAP-146, HPAP-148, HPAP-149, HPAP-155, HPAP-156, HPAP-159, HPAP-160, HPAP-167, HPAP-168, HPAP-172, HPAP-173, HPAP-174, and HPAP-175), were obtained from the PANC-DB portal (https://hpap.pmacs.upenn.edu).

## Acknowledgments

This work was supported by the HIRN Data Scholars Leveraging Multi-Modal Human Islet Research Network Knowledge (LINK) Award (NIH/NIDDK 5U24DK104162-12 to L.W.) and the IIDP Data Resource Trainee Scholar Award (NIH/NIDDK U24DK098085 to L.W.). Z.W. is supported by NIDDK R01DK132651 and R01DK143607. This manuscript used data acquired from the database (https://hpap.pmacs.upenn.edu/) of the Human Pancreas Analysis Program (HPAP; RRID:SCR_016202)^13,23^. The authors thank Xiaozheng Yao, Brian Park, Kai-Ling Chen, and Mitchell J. Machiela for assistance with LDlink concurrent access. LDlink is a publicly available webtool hosted at <u>ldlink.nih.gov</u> by the Integrative Tumor Epidemiology Branch of the Division of Cancer Epidemiology and Genetics and maintained by the Center for Biomedical Informatics and Information Technology. LDlink is funded by the intramural research program of the National Cancer Institute within the US National Institutes of Health.

## Author information

## Author Contributions Statement

L.W. and Z.W. designed the study. L.W. performed all computational analyses. L.W. and Z.W. wrote the manuscript with input from all authors.

